# Extracellular matrix deposition controls early differentiation patterns in human cardiac gastruloids

**DOI:** 10.1101/2025.03.18.643854

**Authors:** Marion F Marchand, Geetika Sahni, Aditya Arora, Selma Serhrouchni, Florian Dilasser, Anaïs Monet, Flora Luciani, Harini Rajendiran, Léa Chabot, Saburnisha Binte Mohamad Raffi, Gianluca Grenci, Jean-Baptiste Sibarita, Rémi Galland, Jin Zhu, Virgile Viasnoff

**Affiliations:** Mechanobiology Institute, National University of Singapore, 5a Engineering drive 1, 117411 Singapore; CINAM, Aix-Marseille Université, UMR 7325 Campus de Luminy 13009 Marseille, France; CENTURI Multi-engineering platform, Campus de Luminy 13009 Marseille, France; IINS, CNRS UMR 5297, Université de Bordeaux 146 Rue Léo Saignat, 33000 Bordeaux, France; BMC2, CNRS IRL3639, 5a Engineering drive 1, 117411 Singapore

**Keywords:** gastruloids, self-organization, extracellular matrix

## Abstract

Amongst the various factors that affect differentiation and tissue organization, the autonomous deposition of extracellular matrix (ECM) has hardly been considered in the context of gastruloids. Using biofunctionalized colloidal particles as artificial organizing centers (aOCs), we patterned differentiation loci within 3D embryonic bodies. Our findings reveal that the spatial distribution of the aOCs induces various pattern organizations, mediated by endogenous deposition of ECM. We investigated how the interplay between meso-endoderm cell epithelial-to-mesenchymal transition, migration into germ layers, and the prolonged maintenance of local stem-cell niches orchestrates gastruloid structure and composition. These factors collectively regulate the complexity of emergent cardiac structures observed at later time points. This work uncovers a critical feedback loop between cell differentiation rates and endogenous ECM deposition patterns, which governs the differentiation paths in scaffolded cardiac gastruloids.

## INTRODUCTION

Gastrulation marks a crucial stage in development, giving rise to the three germ layers – ectoderm, mesoderm and endoderm – while establishing the fundamental body axes of the embryo. Stem cell-derived embryo models, including gastruloids, have recently become valuable tools for investigating these mechanisms *in vitro* ^1–4^. The self-organization properties of gastruloids have been extensively studied, primarily from the perspective of generating an increasingly large variety of cell types, through timed sequences of soluble growth factors treatments. These isotropic stimuli can be complemented by biophysical cues, such as embedding in an extracellular matrix^5^, external scaffolding via micron-scale containers^6,7^, or even 2D adhesive patterns^8–10^. Especially for 3D systems, these cues provide external boundary conditions that guide the development of the gastruloids, leading to spontaneous symmetry breaking and polarized elongation along an anteroposterior axis. Recently, the model has been pushed further towards early organogenesis, with development of heart-like structures^11,12^.

Our current understanding of the self-organization process and lineage specification in organoids/ gastruloids mainly focuses on: *i-* the evolution of cell population transcriptomic states *ii-* the observation of cellular movements that lead to the formation of multicellular structures mimicking aspects of embryonic development. Several studies focus on the role of tissue flows and regulation of cell-cell junctions^13,14^ and cell mechanics^15^ that all contribute to tissue morphogenesis. Although intrinsic ECM deposition is acknowledged as a crucial process shaping morphogenesis during development^16,17^, it remains largely overlooked and poorly understood in gastruloids.

During human gastrulation, mesodermal cells emerge from the epiblast in a region known as the primitive streak, with the Primitive node serving as a key organizing center. As cells ingress through the primitive streak, they undergo an epithelial-to-mesenchymal transition (EMT). This critical process enables them to lose their epithelial characteristics, such as apical-basal polarity and tight cell-cell junctions, and acquire mesenchymal properties, including increased motility and extensive remodelling of the ECM^18^. Cells secrete and interact with ECM components such as fibronectin, laminin and collagens, which in turn provide essential structural support and signaling cues for migration. Cell differentiation, ECM deposition, and cell migration are three interconnected processes that occur on similar timescales during gastrulation, involving both cell-autonomous and collective behaviours. We propose that the intricate balance between these factors dictates tissue organization in gastruloids, ultimately shaping their developmental trajectory and cellular diversity.

To test this hypothesis, we developed an experimental approach to spatially control ESC differentiation. Colloidal particles coated with a combination of growth factors served as artificial organizing centers (aOCs) acting by juxtacrine signaling. These particles were co-assembled with ESCs into embryoid bodies (EBs). We demonstrate that the quantity of beads determines the differentiation rate of epiblast (Epi) cells into meso-endodermal (MesEnd) cells, while their spatial distribution within the EB orchestrates MesEnd and Epi segregation. Moreover, we show that ECM deposition is profoundly influenced by the spatial distribution of aOCs. This ECM modulation governs tissue segregation by altering the ratio of MesEnd microenvironments to Epi microenvironments with consequences on the differentiation states of the cells. Finally, we illustrate how early tissue organization affects subsequent developmental pathways, leading to the formation of diverse complex cardiac structures.

## RESULTS

### Artificial organizing centers control germ layer organization in a 3D gastruloid system

To control the spatial pattern and rate of differentiation of human embryonic stem cells (hESCs) into meso-endoderm cells, we engineered artificial organizing centers (aOCs) using 15 μm Avidin coated polystyrene beads. We treated them with biotinylated heparin to subsequently anchor growth factors (GFs). The beads are then incubated in PBS containing Activin A (100 ng/mL), BMP4 (25 ng/mL), Wnt3a (100 ng/mL) and FGF2 (10 ng/mL) (**Figure 1A** and **Methods**). A subsequent incubation with dilute 1X hESC-qualified Matrigel promotes cell adhesion to the beads. Such a protocol yields individual beads of the size of a single cell onto which stem cells can attach and engage into juxtacrine signaling with all four GFs together. In this study, we used three hESC lines (H1, H9, RUES-GLR). All three lines led to qualitatively similar results. Otherwise noted, the main results showed use H1 cells. We embedded the aOCs in hESCs embryonic bodies according to two protocols (**Figure 1B**). Either we assembled dissociated hESCs and the aOCs in AggreWell-400 plates (StemCell Technologies) to create an isotropic distribution of aOCs in the EB (iso-aOC); or we first centrifuged the aOCs in the AggreWell plates and subsequently seeded the hESCs on top to form EBs with a polarized distribution aOCs (pol-aOC). In both cases, the beads internalized in the EBs with hESCs adhering to them.

**Figure 1:**
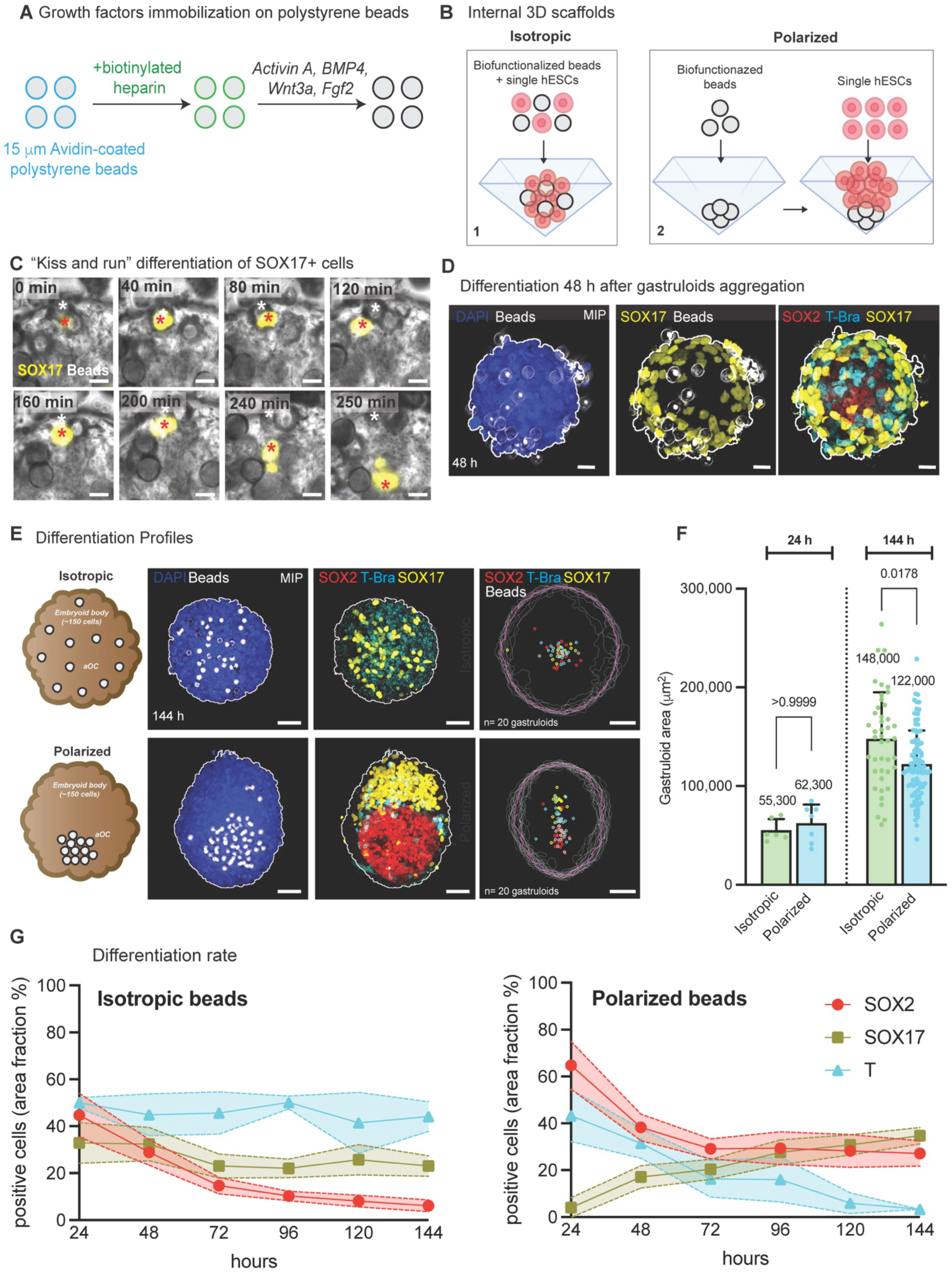
Generation of μ-scaffolded gastruloids using artificial organizing centers. **(A)** Generation of artificial organizing centers by immobilization of a cocktail of growth factors (Activin A, BMP4, Wnt3a, FGF2) on polystyrene beads. **(B)** Experimental procedure to generate isotropic (1) or polarized (2) 3D gastruloids. **(C)** Time-lapse showing a “kiss-and-run” differentiation of a SOX17^+^ endoderm cell (red asterisk) next to a bead (white asterisk). Time is indicated in minutes and t=0 corresponds to the induction of the SOX17 signal in one cell. Scale bar: 10 μm. **(D)** Representative image showing how the differentiation pattern looks like at 48 h after gastruloids aggregation. Scale bar: 20 μm. **(E)** Localization of SOX2, T/Bra and SOX17 transcription factors in isotropic (top row) and polarized (bottom row) gastruloids at 144 h after aggregation. Centroid maps of SOX2, T/Bra, SOX17 and the beads for n=20 isotropic and n=20 localized gastruloids at 144 h after aggregation. Gastruloids shapes (white lines) have been extracted from the DAPI images and an average shape (pink line) is represented. Scale bar: 50 μm. **(F)** Gastruloids size (μm^2^) is showed at 24 h and 144 h after aggregation in iso-aOC and pol-aOC gastruloids. The area was calculated on MIP images of the DAPI channel for each gastruloid (N=1 set of experiment for 24 h, n=6 isotropic gastruloids, n=7 polarized gastruloids; N=6 sets of experiments in isotropic gastruloids at 144 h, n=43 gastruloids; N=9 sets of experiments in polarized gastruloids at 144 h, n=101 gastruloids). Data are represented as mean +/- SD. Statistical comparison made via Kruskall-Wallis test. **(G)** Fraction of positive cells (% of total cells) is showed for each of the germ layer markers SOX2, T/Bra and SOX17 at 24, 48, 72, 96, 120 and 144 h after gastruloids aggregation. This fraction corresponds to the area occupied by each marker divided by the area occupied by DAPI for each image. 2 sets of experiments (1 experiment performed with H1 cells and 1 experiment with RUES2-GLR cells) were pooled and n=6-45 gastruloids were analyzed at each timepoint for each marker. Data displayed on graph corresponds to the mean and error with a 95% confidence interval for each timepoint.

We first tested the ability of aOCs to induce cell differentiation. We used the RUES-GLR cell line, expressing a SOX17-tdTom reporter to follow the individual dynamics of endoderm differentiation and the consequential epithelial-to-mesenchymal transition (EMT). 24 hours after aggregation (haa), we transferred the EBs into micro-fabricated wells designed for single-objective light-sheet microscopy (soSPIM)^19,20^ and performed live imaging to follow the differentiation dynamics of the hESCs over 3 days. We observed an induction period of 36–48 hours before the initial expression of the transcription factor SOX17, comparable to induction by soluble growth factors (**Supplementary Figure 1A**). However, in the case of aOCs, differentiation was restricted to the immediate vicinity of the aOCs (**Figure 1C**). The differentiating cell remained in close contact with the aOC for approximately 3 hours before migrating away from the aOC, suggesting and EMT process. Some aOCs were surrounded by more than one SOX17^+^ cell, while others had none. This variability may stem from the ∼1 day heterogeneity induction time following initial contact with the beads. This kiss-and-run mechanism (**Video 1**) operates independently of the spatial distribution of aOCs and triggers the differentiation of multiple SOX17^+^ cells within EBs. This process is obvious in gastruloids at 48 haa (**Figure 1D**), where several SOX17^+^ cells are distributed within the EB. After 72h, the aOCs lose their differentiation capability likely due to the degradation of the GFs on their surface. However, EBs continue to grow up to 144 h after aggregation, reaching 250-300 μm in size (**Video 2** and **Figure 1E**).

We identified day 6 (144 haa) as a turn-point in the development of the gastruloids. At this stage, each aOC distribution led to a stark difference in tissue organization. Immunostaining and confocal imaging revealed a homogeneous distribution of T/Brachyury and SOX17 positive cells and the absence of SOX2^+^ cells (epiblast marker) for iso-aOC (**Figure 1E**) which is comparable to a treatment with soluble GF (**Supplementary Figure 1A**). In this case, we observed no symmetry breaking (i.egatruloid elongation). By contrast, pol-aOCs gastruloids organized in two distinct poles, one containing the SOX2^+^ cells and the second SOX17^+^ and T/Bra^+^ cells. We validated the robustness of the stereotypic arrangement of cell populations in both iso-aOCs and pol-aOCs by quantifying the average spatial distribution of various germ layer markers across 20 randomly selected gastruloids (**Figure 1E and Methods**). In the SOX2^+^ pole, the cells retained other pluripotency markers such as OCT4 and NANOG unless they were in direct contact with the aOCs, demonstrating the localized nature of the differentiation (**Supplementary Figure 1B**). The SOX17^+^ pole included FOXA2^+^ and CDX2^+^ cells, confirming their meso-endodermal lineage (**Supplementary Figure 1C**). The relative cell proportion between the SOX17^+^ pole and SOX2^+^ pole varied linearly with the number of aOCs, provided it exceeded a threshold of 10-12 beads, below which no differentiation occurred (**Supplementary Figure 1D**). Immunostaining of E- and N-cadherins revealed that SOX2^+^ cells are also E-cadherin positive, whereas SOX17^+^ cells are N-cadherin positive, confirming that the cadherin expression profile follows the EMT expression pattern (**Supplementary Figure 1E**).

We then compared the dynamics of cell differentiation for both cases using a time-resolved series of immunostainings on the same batch of gastruloids (**Supplementary Figure 1F**). The overall growth of the gastruloids was similar in both cases, leading to gastruloids of similar sizes at 144 haa (**Figure 1F**). However, the SOX2^+^ population decreased overtime between day 1 to day 4 (96 haa) in the iso-aOCs gastruloids and was replaced by a constant population of SOX17^+^ and T/Bra^+^ cells (**Figure 1G**). By contrast, for pol-aOC, the SOX2^+^ population decreased until it reached a plateau at 72 haa, with the SOX17^+^ population becoming gradually dominant over the T/Bra^+^ cells (**Figure 1G**). Together, these results strongly suggest that both the localization and number of aOCs govern the dynamics of cell differentiation, enabling a controlled balance of cell types in gastruloids and leading to distinct tissue organizations.

### Endogenous extracellular matrix deposition guides tissue formation in polarized gastruloids

Cell sorting during EMT is usually explained based on differential adhesion/tension between cells. The observation that the spatial distribution of aOCs led to different tissue organization and cellular composition could not be explained within this framework. We therefore explored an alternative explanation centered on the dynamic local formation of the cellular microniche, driven by the secretion of endogenous extracellular matrix, which could regulate both migration and direct cell differentiation.

We hypothesized that the enhanced differentiation of SOX2^+^ cells in the iso-aOC case was related to a local disruption of the microniche that ensures their maintenance. We thus performed single-cell RNA sequencing (sc-RNA seq) on several pools of polarized gastruloids at 144 haa (**Supplementary Figure 2A**). A thorough analysis of the specificity of expression of ECM genes revealed that fibronectin (*FN1*) was one of the main differentially expressed genes between meso-endoderm cells and epiblast cells, as compared to other ECM genes such as *COL4A1*, *COL4A2* and *LAMB1* (**Supplementary Figure 2A**). We thus investigated its role in maintaining the homeostasis of local microniches. An intricate fibrillar FN meshwork was observed within the meso-endoderm regions of both iso-aOC and pol-aOC gastruloids (**Figure 2A**). Additionally, 81% of pol-aOC gastruloids exhibited thick FN bundles extending across the pluripotent pole, linking the vicinity of the aOC to the meso-endoderm pole. (**Figure 2A**, **Supplementary Figure 2B-C**). The epiblast pole presented scattered SOX17^+^ cells that colocalized in 94% of the cases with the FN bundles **(Supplementary Figure 2C**). Such bundle organization was never observed in iso-aOC gastruloids, suggesting that the clustering of aOCs generated FN tracks capable of guiding cell migration while reinforcing themselves through repeated localized FN secretion. To further support this hypothesis, we selectively tested the affinity of SOX17^+^ and SOX2^+^ cells for different ECM proteins using cell spreading and cell migration assays. H1 cells were either maintained as stem cells or differentiated into SOX17^+^ cells before being plated on top of different ECM substrates including fibronectin (**Figure 2C**). Pluripotent H1 cells moderately adhered to fibronectin-coated substrates but could not migrate on them (**Figure 2D**). By contrast, SOX17^+^ cells adhered twice better on fibronectin and their migration was significatively enhanced (**Figure 2D**). For SOX17^+^ cells, we also tested other ECM substrates including collagen I, laminin and Matrigel (**Supplementary Figure 2E-F**). Fibronectin was the only substrate providing strong adhesion, as well as fast and efficient migration (**Supplementary Figure 2E-F**). These data further support the idea that SOX17^+^ cells follow the fibronectin they secrete, as previously suggested in the “motility-based deposition” model of fibronectin^21,22^. The secreted fibronectin may serve as a ‘memory’ for migrating cells, with the tracks guiding the movement of differentiated cells toward a single pole.

**Figure 2:**
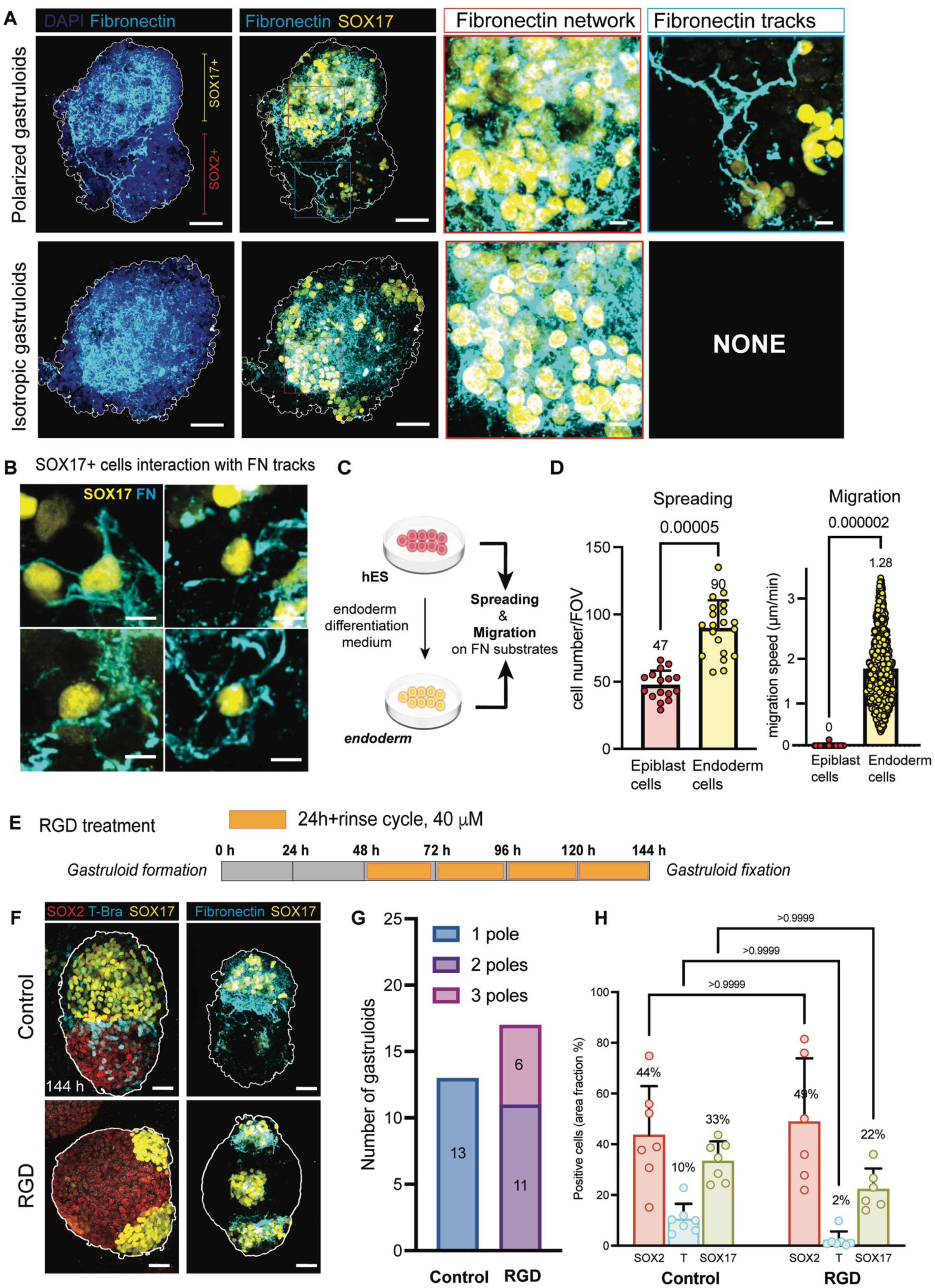
Fibronectin deposition in the form of tracks in polarized gastruloids. **(A)** Deposition of fibronectin in polarized (top row) and isotropic (bottom row) gastruloids at 144 h after aggregation. A dense network of fibronectin (red inserts) is present in both isotropic and polarized gastruloids in the SOX17^+^ region. Some fibronectin long fibers named as tracks (blue inserts) are present only in the polarized gastruloids. Scale bars: 50 μm; zooms: 10 μm. **(B)** Examples of fibronectin tracks associated with SOX17^+^ cells. Scale bar: 10 μm. **(C)** Experimental procedure for 2D experiments; hES were first differentiated into endoderm cells expressing SOX17 or kept as undifferentiated stem cells before detachment and spreading on fibronectin coating. Cell adhesion and migration was then assessed. **(D)** Number of cells/fields of view was assessed after 4 h spreading on fibronectin. 20 fields of view were analyzed for each condition. Statistical comparison made via Kruskall-Wallis test. Cell migration speed (μm/min) was analyzed on brightfield time-lapse movies, 1 image every 5 minutes for 4 hours. Sample size is n=1379 analyzed tracks for endoderm cells plated on fibronectin. **(E)** Schematic representation of RGD treatment experiments. **(F)** Localization of SOX2, T/Bra and SOX17 markers as well as fibronectin deposition (cyan) in control and RGD-treated gastruloids at 144 h after aggregation. Scale bar: 50 μm. **(G)** Number of SOX17^+^ poles in control and RGD-treated gastruloids at 144 h (n= 13 control gastruloids; n=17 RGD-treated gastruloids). **(H)** Fraction of positive area (% of total area) is represented for each of the SOX2, T/Bra and SOX17 markers in control (left) and RGD-treated (right) gastruloids at 144 h. n=6-7 gastruloids analyzed for each condition. Data are represented as mean +/- SD. Statistical comparison made via Kruskall-Wallis test, ns indicates p>0.9999.

We then antagonized the cellular interactions with FN by adding RGD peptides to the culture media between 48 and 144 haa (**Figure 2E**). Soluble RGD peptides compete with RGD-containing ECM proteins such as FN and laminin to bind to β subunits of integrins. We started treating the cells at 48 haa, corresponding to the onset time of differentiation, according to our live imaging experiments (**Supplementary Figure 1A** and **Video 2**) until 144 haa. Pol-aOCs gastruloids treated with RGD organized multiple SOX17^+^ cells poles around the SOX2^+^ pole (**Figure 2F-H**). Each of these poles presented a dense fibrillar meshwork of fibronectin (**Figure 2F**). Fibronectin tracks were hardly detected upon RGD treatment (**Figure 2F**). Together, these data identify FN bundles as migratory tracks that facilitate their own local deposition by meso-endoderm cells through enhanced collective migration. These tracks emerge from the spatially restricted region where H1 cells differentiate and undergo EMT and do not form when differentiation occurs isotropically. They play a key role in organizing cells into distinct poles.

Fibronectin has been reported to foster the differentiation of pluripotent stem cells towards the mesodermal fate, with a specific bias toward precardiac mesoderm^23,24^. We therefore hypothesized that a localized imbalance between FN deposition and collagen would promote the differentiation of epiblast cells into mesoderm. We reasoned that in iso-aOCs, the random deposition of FN could disrupt the local microniche, leading to the complete differentiation of epiblast cells into meso-endoderm. In contrast, in pol-aOCs, FN tracks would guide FN deposition during meso-endoderm cell migration, preserving local microniches that support pluripotency maintenance. We hence tested this hypothesis probing the distribution of collagen IV and laminin, the main components of the basement membrane. They were ubiquitously expressed by all cell types present in the polarized gastruloids (**Supplementary Figure 2A**). We thus checked for their localization in 144 haa gastruloids after immunostainings. In iso-aOCs, they formed a diffuse discontinuous network and were present homogeneously (**Figure 3A** and **Supplementary Figure 3A-B**). However, in pol-aOCs gastruloids, laminin and collagen IV were additionally present in a basement membrane sheet-like structure, at the border between the SOX2^+^ and the SOX17^+^ areas (**Figure 3A** and **3B**). We observed a stereotypical alignment of SOX17^+^ cells along the BM (**Figure 3B**), as well as an apico-basal polarization of epiblast cells at the border (**Supplementary Figure 3C**). The BM was discontinuous with signs of cells crossing in the cracks (**Figure 3B**), reminiscent of the formation of basement membranes between the different germ layers in mouse embryos^25^. The quantification of laminin deposition indicated that the total amount of deposited laminin was similar in both cases (**Supplementary Figure 3A-B**). However, laminin localization was different between iso-aOC and pol-aOC gastruloids, suggesting that its secretion is not impaired whereas its remodeling and localization are. A targeted pulse digestion of the basement membrane using a collagenase burst at 96 or 120 haa (i.e., after gastruloid polarization) led to a clear separation between the pluripotent and meso-endoderm poles (**Supplementary Figure 3D-E**). These findings suggest that collagens play a crucial role in maintaining the cohesion of the two poles following the final polarization of pol-aOC gastruloids.

**Figure 3:**
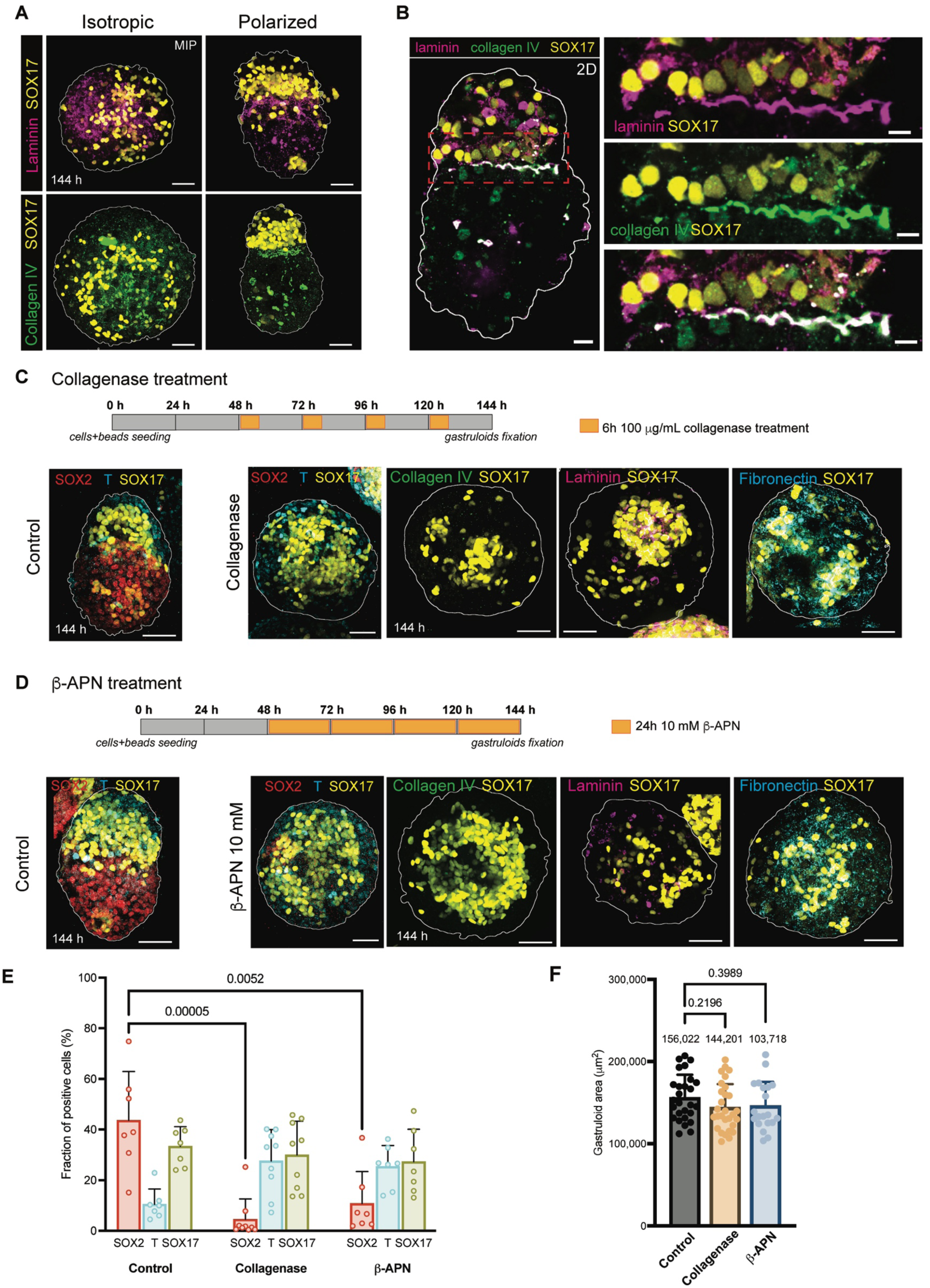
Collagen IV deposition and maintenance of the stem cell niche in polarized gastruloids. **(A)** Deposition of laminin (magenta, top row) and collagen IV (green, bottom row) in iso-aOC and pol-aOC gastruloids at 144 h. Scale bars: 50 μm. **(B)** Basement membrane deposition and cell orientation in pol-aOC gastruloids, showing collagen IV and laminin colocalizing at the interface between the SOX17^+^ and SOX2^+^ areas. Scale bar: 50 μm, zoom: 10 μm. **(C)** Experimental outline of the protocol used for collagenase treatment of gastruloids. Collagenase (100 μg/mL) was added to cell culture medium every day from 48 h and incubated during 6 h before removing it from the culture medium. Samples were fixed at 144 h. Localization of SOX2, T/Bra and SOX17 in control and collagenase-treated gastruloids at 144 h. Scale bar: 50 μm. ECM deposition in gastruloids treated with collagenase from 48 h for 6 h every day is also shown. Collagen IV, laminin and fibronectin deposition are showed at 144 h. Scale bar: 50 μm. **(D)** Experimental outline of the protocol used for β-APN treatment of gastruloids. 10 mM β -APN was added every 24 haa from 48 haa until gastruloids fixation at 144 haa. Localization of SOX2, T/Bra and SOX17 in control and b-APN-treated gastruloids at 144 h. Scale bar: 50 μm. ECM deposition in gastruloids treated with b-APN fixed at 144 h. Scale bar: 50 μm. **(E)** Fraction of positive area/total area (%) for SOX2, T/Bra and SOX17 markers in control (n=7 gastruloids), collagenase (n=9 gastruloids) and β -APN (n=7 gastruloids)-treated gastruloids at 144 haa. Data are represented as mean +/- SD. Statistical comparison made via Kruskall-Wallis test. (**F**) Size (area in μm^2^) of control (n=27 gastruloids), collagenase (n=29 gastruloids) and β -APN (n=20 gastruloids)- treated gastruloids at 144 haa. Data are represented as mean +/- SD. Statistical comparison made via one-way ANOVA (p= 0.2196 and p=0.3989).

We then applied a daily pulse disruption of collagens (6 hours per day) starting at 48 haa using collagenase burst digestion, which led to a significant reduction in the SOX2^+^ population (**Figure 3C** and **3E**). A similar phenotype was observed after treating cells with 10 mM β-APN, a LOX inhibitor, impairing collagens cross-linking (**Figure 3D-E**). We showed that β-APN treatment was dose-dependent, increasing the defects of SOX17 spatial patterning when used at 10 mM instead of 1 mM (**Supplementary Figure 3F**). However, treating cells with β-APN from 96 or 120 h led to no significant change in gastruloids polarization (**Supplementary Figure 3G**), showing that collagens cross-linking is crucial during the first 72 h, when most of the SOX17 cells are being differentiated. For both treatments, the gastruloids retained a growth rate identical to controls (**Figure 3F**) and no abnormal cell death was observed in phase contrast microscopy. We concluded that the loss of expression of SOX2 upon collagenase and β-APN treatments resulted from an enhanced differentiation and not from the death of the SOX2^+^ cells. At 144 haa, the treated pol-aOC gastruloids were phenotypically indistinguishable from the control iso-aOC gastruloids (**Figure 3C-D**), with fibronectin fibers uniformly distributed throughout the entire structure. Our results strongly suggest that the balance between FN and collagen is essential for maintaining a pool of pluripotent cells in gastruloids, offering a coherent explanation for the differences observed between iso-aOC and pol-aOC gastruloids. We then investigated the downstream impact of prolonged epiblast cell maintenance on cellular diversity.

### Cellular diversity differs between pol-aOCs and iso-aOCs

We performed single-cell RNA sequencing (sc-RNA seq) on 48 haa and 144 haa on pol-aOC gastruloids using 10X Genomics Chromium to explore the diversity of cellular composition obtained over time. After quality control, during which doublets and low-quality cells were filtered out, we performed Uniform Manifold Approximation and Projection (UMAP) and identified 8 cell clusters based on differentially expressed genes (**Figure 4A**). We annotated these clusters based on combination of putative lineage markers, as established by previous single-cell transcriptomics studies of gastrulating human embryos^26^. The comparative UMAP of pol-aOC gastruloids at 48 and 144 haa reveals that at 48 haa, cells primarily belonged to epiblast, nascent mesoderm, endoderm, and ectoderm lineages. At 144 haa, the gastruloids matured while preserving an epiblast cluster. They showed more cellular diversity, including emergent mesoderm cells, advanced mesoderm cells, hematopoietic and endothelial progenitors, showing that the mesoderm cluster has significantly increased and matured. They also showed an increase in the amount of ectoderm cells, as well as neural progenitors cells. We detected neural crest cells progenitors for central nervous system (*POU3F2+* or Oct7) as well as peripheral nervous system (*POU3F1+* or Oct6) at 144 haa (**Supplementary Figure 4A**). The overall cell population splits between epiblast SOX2^+^ cells, and cardiac mesoderm cells, which co-express *HAND1* and *HAND2*, two master genes regulating heart morphogenesis and development (**Figure 4B**). This suggests that the mesoderm pole is predominantly cardiac, aligning with the pattern of FN deposition.

**Figure 4:**
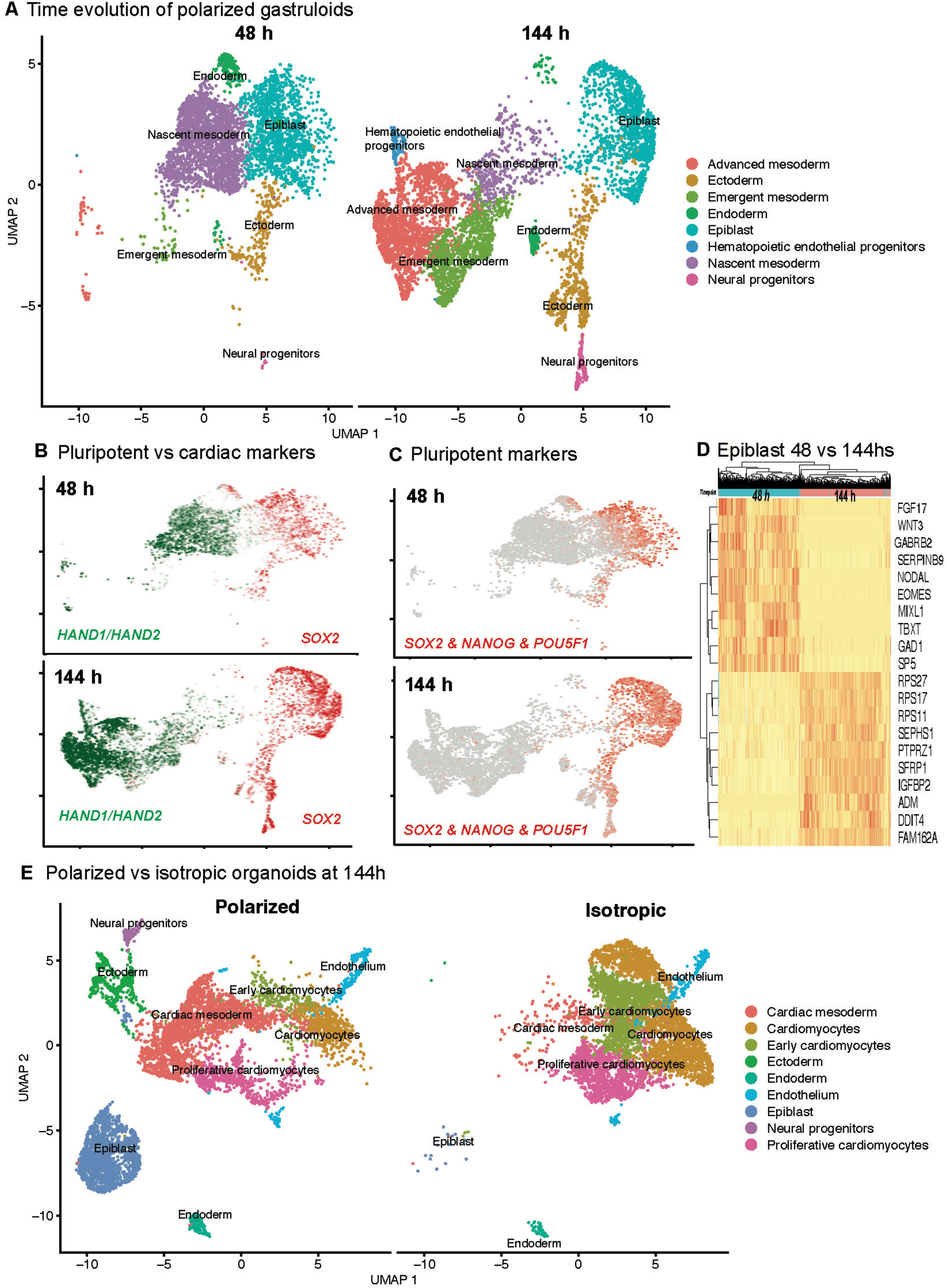
Single-cell RNA sequencing of isotropic and polarized gastruloids **(A)** UMAP plot representation showing the projection of 48 haa and 144 haa gastruloids, clusters have been obtained from 10,709 isolated cells. (**B**) In green, the expression of both *HAND1* and *HAND2* is shown and in red the expression of *SOX2*, showing two distinct populations. **(C)** UMAP plots showing the combined expression of pluripotent markers *SOX2*, *NANOG* and *POU5F1* in 48 and 144 haa gastruloids. (**D**) Heatmap showing the top differentially expressed genes in the epiblast clusters in 48 and 144 haa gastruloids. **(E)** UMAPs plots showing clusters obtained from 12,622 cells isolated from 144 haa polarized or isotropic gastruloids.

At 48 haa, the epiblast cluster expresses high levels of *FGF17*, *FGF4*, *WNT3*, *NODAL*, *EOMES*, *TBXT*, *MIXL1* (**Figure 4D** and **Supplementary Figure 4B**). All these genes are involved in maintaining the epiblast state, as well as driving mesodermal and primitive streak differentiation. At 144 haa, the epiblast cells are no longer *bona fide* epiblast cells like at 48 h. They still expressed the pluripotency markers *SOX2*, *POU5F1* and *NANOG*, (**Figure 4C**), but they are in the process of committing to neural/ectodermal lineages both for PNS and CNS, as demonstrated by the comparison of these cells at 48 and 144 haa (**Figure 4D**). They express high levels of ribosomal protein genes, suggesting that this population is in a state of active protein synthesis and cell growth. They also express *PTPRZ1*, indicating a neural progenitor identity, and *SFRP1*, responsible for Wnt signaling inhibition, which could indicate a shift towards neural differentiation for these cells. The pol-aOC gastruloids hence evolved towards cardiac lineages showing cardiac and vascular progenitors, while maintaining an endodermal pool of cells, and developing a neural lineage. A portion of the cellular diversity at 144 haa consists of cells that retain a high degree of stemness.

We then compared the transcriptomes of pol-aOC to iso-aOC gastruloids at 144 haa (**Figure 4E**). In iso-aOC gastruloids, the epiblast population is no longer present, consistently with our immunostaining data. The ectoderm and neural progenitor clusters were also significantly reduced, while the endoderm cluster was maintained but slightly decreased, as also indicated by FOXA2 expression (**Supplementary Figure 4C**). In the iso-aOC condition, we observed more mature mesoderm lineages, including a larger cluster of proliferative cardiomyocytes in the G2/M phase, characterized by high expression of *NEK2* and *PLK1*, two key cell cycle regulators. They also showed a larger cluster of late cardiomyocytes cells, expressing high levels of *MYH6*, *SLC8A1*, *UNC45B*, involved in cardiac muscle contraction, calcium homeostasis and sarcomere organization. The hematopoietic and endothelial progenitors cluster remained similar in iso-aOC and pol-aOC gastruloids. Pol-aOC gastruloids highly expressed early mesoderm markers, corresponding to the nascent and emergent mesoderm clusters, including *EOMES*, *MIXL1*, *TBX6* and *MESP1* (**Supplementary Figure 4D**). The data thus suggest that the prolonged maintenance of epiblast cells in pol-aOC gastruloids leads to a slower maturation of the cardiac lineages while favoring neural progenitors and ectoderm. In contrast, the iso-aOC gastruloids reached a less diverse but more mature cardiac phenotype, with most of the cell population expressing some cardiomyocytes gene markers such as *TNNT2*.

### Enhanced the complexity of cardiac structures in day 12 in pol-aOC gastruloids

We then focused on cardiac mesoderm development. **Figure 5A** shows the spatial organization of HAND1-expressing cells, a transcription factor crucial for primary heart field formation and cardiac morphogenesis. HAND1 expression was detected as early as 48 haa in both iso-aOC and pol-aOC gastruloids (**Figure 5A**). However, in iso-aOC gastruloids, HAND1^+^ cells remained scattered throughout the structure until day 12, whereas in pol-aOC gastruloids, they predominantly localized within the mesoderm pole and formed a monolayer surrounding the epiblast pole by day 4. At day 12, both types of gastruloids reached similar volumes (**Figure 5B)**. The iso-aOC remained spherical whereas the pol-aOC displayed a pronounced elongated shape (**Figure 5B**). Only the pol-aOC gastruloids developed a pulsatile behavior at one of their poles (**Figure 5C** and **Video 3**). Particle Imaging Velocimetry and kymograph generation revealed two distinct dynamics. The capping regions collectively contracted over the whole length of the gastruloids, beating at around 0.5 Hz, in line with previous reports on early cardiomyocytes^27^. The internal connected regions show erratic vibrations around 10 Hz (**Figure 5D**) that could possibly be explained by the interaction with neural crest-derived autonomic neurons, interacting with the adjacent cardiomyocytes. By contrast, iso-aOC gastruloids showed no sign of any contractile activity.

**Figure 5:**
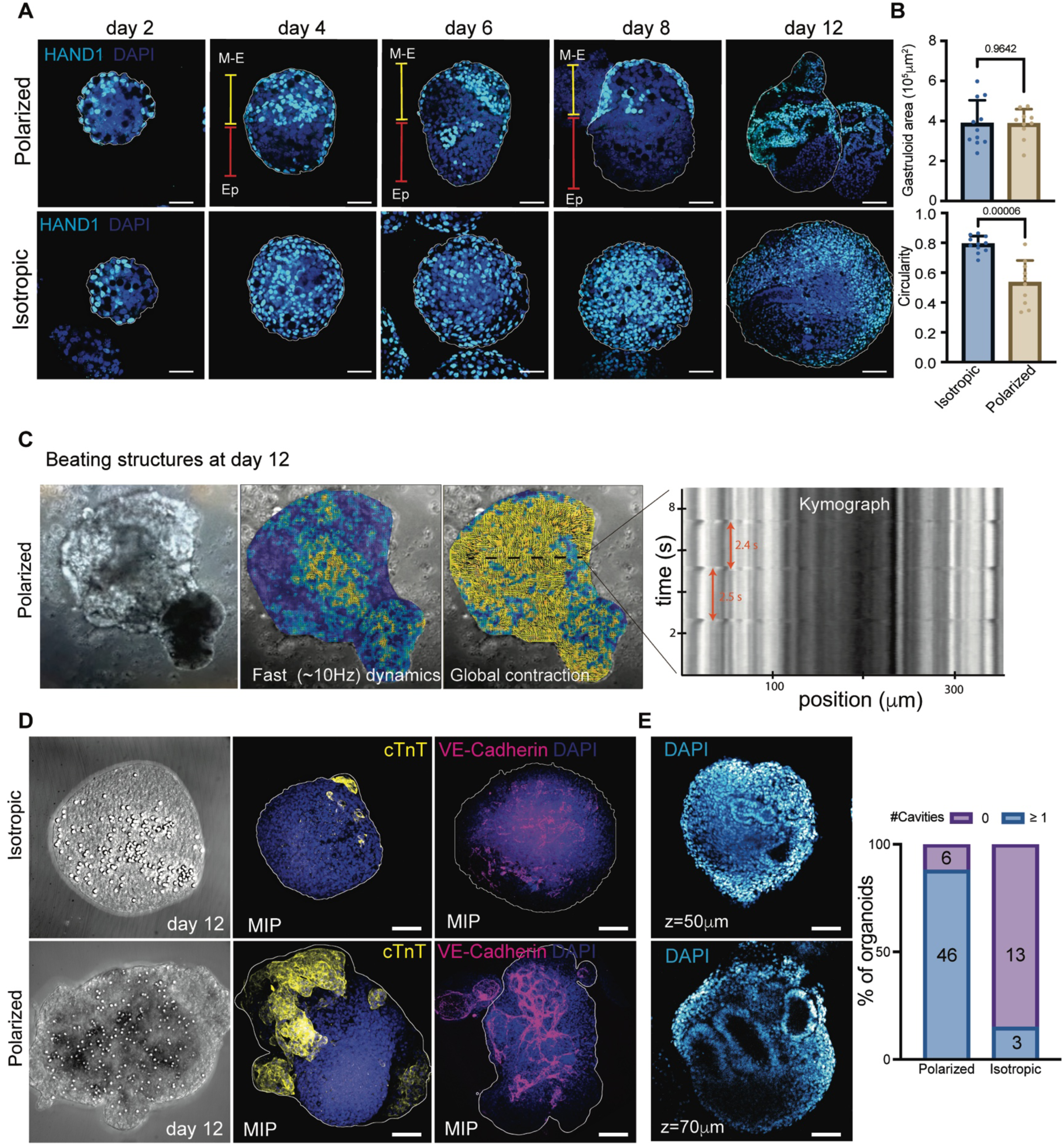
Cardiac lineage development and emergence of cardiovascular structures in polarized gastruloids. **(A)** Localization of HAND1 in pol-aOC and iso-aOC gastruloids at 48, 96, 144, 192 haa (scale bar: 50 μm) as well as after 12 days of culture (scale bar: 100 μm). Ep= epiblast; M-E= meso-endoderm. **(B)** Gastruloid area and circularity of isotropic (n=11) and polarized (n=10) gastruloids at day 12 is also shown. Data is shown as mean +/- SD. Statistical analysis made via unpaired t-tests. **(C)** PIV analysis of the beating structures in polarized gastruloids at day 12 (also see **Video 4**) showing fast contractions dynamics (10 Hz) as well as global slow contractions. A kymograph is shown to illustrate the global contractions. **(D)** Expression of cardiac troponin T (cTnT, yellow) and VE-Cadherin (magenta) in isotropic and polarized gastruloids at day 12. Scale bar: 100 μm. **(E)** DAPI staining is shown for 1 slice of the gastruloids to highlight the presence of cavities inside the gastruloids. Quantification of the number of cavities (0 or > 1) is shown in pol-aOC (n=52) and iso-aOC (n=16) gastruloids.

Contracting regions correlated with the expression of cardiac troponin T (cTnT), a protein that modulates cardiac muscle contraction through its interactions with neighboring thin filament proteins (**Figure 5D**). Iso-aOC gastruloids displayed very small clusters of cTnT^+^ cells that were not organized or connected. VE-cadherin was expressed in both conditions, but the vessel-like structures formed in the pol-aOC gastruloids were highly connected and formed a large vascular network expanding throughout the whole gastruloid (**Figure 5D**). 90% (n=52 gastruloids) of pol-aOC gastruloids, developed large internal (>100μm) cavities that were only detected in 15% of the iso-aOC (n=26 gastruloids) (**Figure 5E**). Human heart morphogenesis involves a series of developmental events leading to the formation of a multi-chambered organ, the formation of cavities within the developing heart is thus a crucial process and cavitation is necessary to form the endocardial lumen^28^. We found cavities with different morphologies as exemplified in **Figure 6A**. We checked for the presence of endocardium markers and noticed several types of cavities were involving SOX17^+^ cells (**Figure 6B**). Out of a pool of 27 pol-aOC gastruloids, 66% (n=27) showed type 1 cavities, i.e. small lumen of 100-150 μm lined by SOX17^+^ cells and surrounded by fibronectin; 30% had very large cavities (400-500 μm) still lined by a multi-layer sheet of SOX17^+^ cells surrounded by fibronectin. This structure could correspond to a more mature stage of endocardium development; 37% had long vessel-like organization of SOX17^+^ cells, with thin and elongated nuclei aligned along specific axes (**Figure 6B**). Endocardium contributes to trabecular vessel formation through endocardial-to-mesenchymal transition^29^. The above structures are reminiscent of such a process in pol-aOC gastruloids. We also observed cavities formed by SOX2^+^ cells surrounded by a layer of collagen IV (**Figure 6C**). These structures could originate from ectoderm/neural progenitors cells that were previously detected in the scRNA-seq dataset (**Figure 4D**). These cells could support heart formation, as we know that neural crest cells interact with the heart fields to influence early cardiac morphogenesis^30^. Finally, we observed other types of cavities in which the polarity was inverted, i.e. lined internally with fibronectin (**Figure 6D**) that could correspond to the formation of valve primordia in the developing heart^31^.

**Figure 6:**
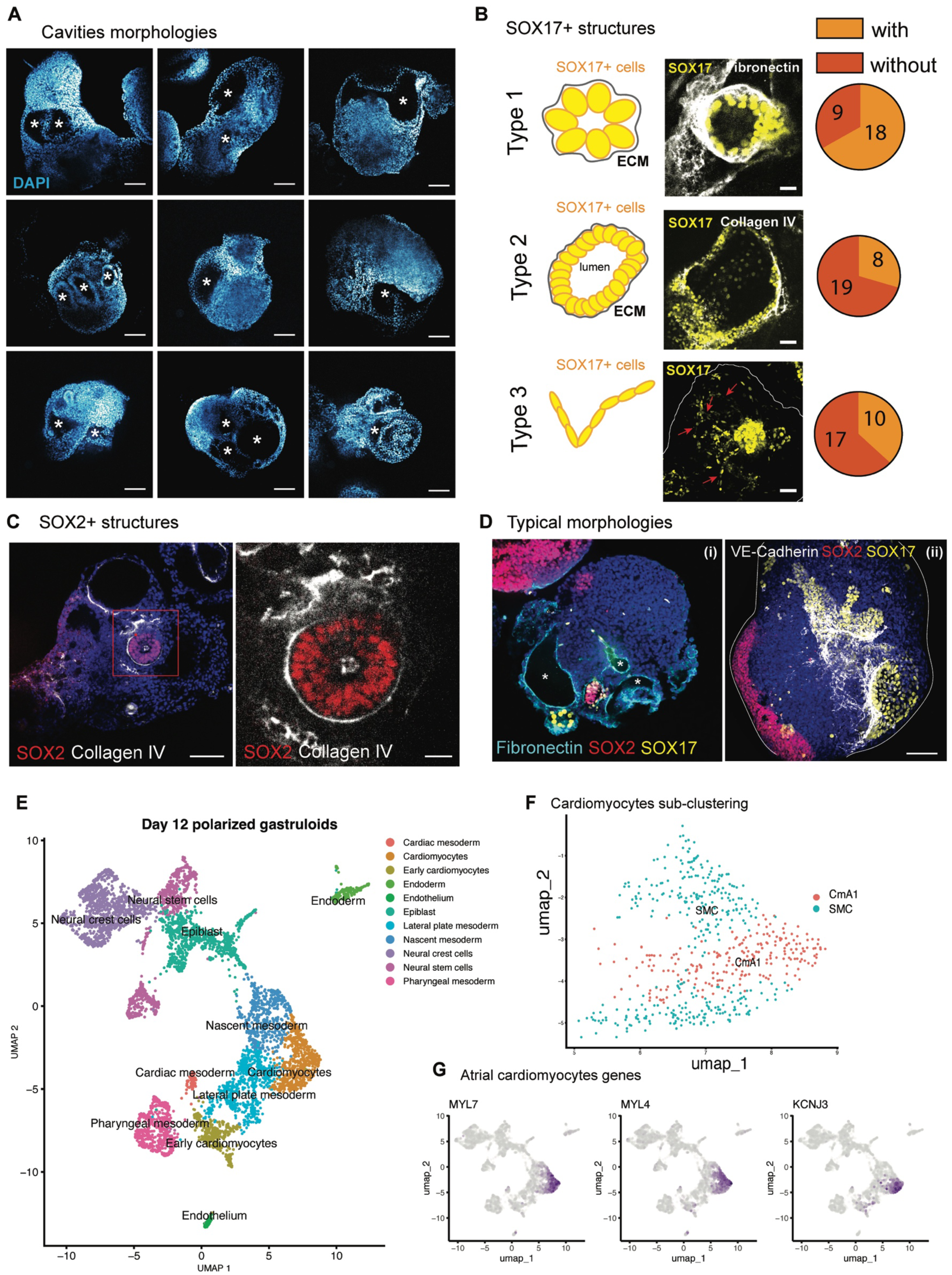
Cardiovascular features in polarized gastruloids at day 12. **(A)** Single plane images of DAPI stainings in 9 different polarized gastruloids at day 12, showing different morphologies of cavities inside (white asterisk). Scale bar: 100 μm. **(B)** Endocardium morphogenesis in polarized gastruloids at day 12. Different types of SOX17+ cavities are observed, depending on the size (type 1 or 2). Some vessels-like structures are also present (type 3, red arrows). Scale bar: 50 μm. For each type of SOX17+ structure, a quantification has been done to decipher whether they are present or not in n=27 gastruloids. **(C)** Illustration of SOX2+ structures (red) ensheathed in a collagen IV layer (grey) Scale bar: 100 μm, zoom: 20 μm. **(D)** Typical morphologies of (i) cavities displaying an internal lining of fibronectin (white asterisks) and (ii) vessel-like structures positive for VE-cadherin supported by SOX17+ cells areas. Scale bar: 100 μm. **(E)** UMAP plot showing clusters obtained from 5,532 cells extracted from pol-aOC gastruloids at day 12. **(F)** UMAP plot showing sub-clustering of the “cardiomyocytes” cluster, divided into Cardiomyocytes A1 and smooth muscle cells. **(G)** UMAP plots showing the expression of atrial cardiomyocytes genes markers in day 12 gastruloids.

sc-RNA sequencing of pol-aOC gastruloids at day 12 confirmed the broad cellular diversity previously inferred from immunostainings (**Figure 6E**). Based on differential gene expression and using the CASSIA annotation tool for single-cell RNA-sequencing data^32^, we identified 11 different clusters. The overall population divides into two main cell identities corresponding to neural and cardiac lineages. The cardiomyocyte cluster predominantly expresses genes associated with atrial identity, including *MYL7*, *KCNJ3*, *MYL4* (**Figure 6G**). In contrast, genes linked to ventricular cardiomyocyte identity, such as *MYL2*, *MYH7*, *IRX4* show lower expression levels. These findings suggest that the gastruloids exhibit a predominantly atrial-like identity. We sub-clustered this group and re-annotated it using an additional database derived from scRNAseq data of human fetal hearts^33^. Sub-clustering revealed a dual population of cardiomyocytes and smooth muscle cells (**Figure 6F**) expressing *ACTA2*, *TAGLN*, and *CNN1* (**Supplementary Figure 5C**). Finally, we examined the expression of endocardial markers and found that, in line with our stainings, the endothelial cell cluster strongly expresses genes associated with endocardial identity, including *PECAM1*, *KDR*, *CDH5*, and *NFATC1* (**Supplementary Figure 5C**). We further identified a pharyngeal mesoderm cluster, which is typically associated with development of cardiac structures and the craniofacial musculature^34,35^. These cells express high levels of *NR2F2*, *FOXP2*, *EYA1*, *SIX1* and *DLX1*, all involved in craniofacial mesoderm and pharyngeal development. Altogether, our data thus provide evidence that we developed a 3D gastruloid model exhibiting both neural and cardiac features. Notably, the cardiomyocytes population within this model shows a distinct atrial-like identity, reflecting the early stages of heart morphogenesis. This model recapitulates key aspects of cardiac development, including emergence of cardiomyocytes subtypes and the formation of functional tissue architecture.

## DISCUSSION

Our study demonstrates that artificial organizing centers can guide the differentiation of gastruloids based on their spatial distribution. The intricate development of gastruloids into cardiac structures over 12 days occurred without the need for soluble growth factors or external extracellular matrix. Instead, their self-organization was driven by aOCs, which served as initial differentiation loci for pluripotent stem cells at controllable rates. The changes in cell diversity we observed originates from the varying commitment rates of pluripotent stem cells to different germ layers. While differentiation initially occurs near the aOCs, it is further regulated downstream by the deposition of fibronectin, endogenously secreted by meso-endoderm cells.

Our data suggest that the local ECM composition influences pluripotent stem cell differentiation, consistent with findings in more mature stem cells, such as mesenchymal and neural stem cells^36^. A higher concentration of fibronectin biases differentiation toward cardiac mesoderm, while a local microniche rich in collagens supports stem cell maintenance, as previously demonstrated with collagen I^37^. One of the key findings of this study is that ECM deposition plays a crucial role in regulating and patterning gastruloids, and likely other organoid types as well. *In vivo*, ECM components provide structural support and biochemical signals that guide lineage segregation and cellular movements, mirroring their functions during embryogenesis. Previous data already evidenced that ECM remodeling plays a key role during embryonic development. More specifically, ECM gradients have been shown to regulate cell differentiation and patterning in mouse embryos^38,39^ but also in drosophila embryos^40,41^. We argue that in the context of organoids, much focus has been granted to cells, overlooking the importance of endogenous ECM deposition.

A second realization that our data support is that the spatial localization of aOC can turn the cell-autonomous process of ECM secretion into a collective effect by creating large FN bundles serving as migratory tracks. We propose that these tracks form by a self-reinforcing mechanism of meso-endoderm cells channeling through preferred migratory paths while secreting FN. The formation of tracks has a dual effect. First, they funnel the migration from the aOC pole towards a single meso-endoderm pole. Second, they locally concentrate FN deposition, hence preserving the collagen niches necessary to maintain the stem cells. Our data thus advocate for a positive feedback loop between cell migration and fibronectin deposition, that compete with the collagen-rich micro-environment needed for epiblast cells maintenance/differentiation. Supporting our results, fibronectin has already been shown to guide neural crest cells during their migration in the chick embryo, creating a scaffold for trailing cells^21^. Fibronectin is also a substrate for meso-endodermal precursors cell migration during the gastrulation process^42^, further supporting the idea that the FN tracks observed in our model can direct the migration of differentiated cells toward a single pole. Finally, it is well known that fibronectin can bind to several growth factors^43^ and could thus act as a platform for the binding and presentation of signaling cues to the migrating cells.

The aOC approach presents a compelling alternative to soluble growth factor treatments, potentially offering greater spatial control over differentiation. Its application to other organoid systems is currently being explored. Notably, this method provides a trackable means of inducing differentiation without compromising the subsequent complex development of gastruloids into fetal cardiac organoids. While a full characterization of cardiac organization remains for future work, the observed cellular complexity aligns well with previously reported cardiac gastruloids and cardioids ^11,12,44,45^. Our cardiac gastruloid model acquired large-scale beating phenotypes over 12 days of differentiation. While previous models have demonstrated the formation of heart tube-like structures^11^ or multi-chambered heart-like structures^46^, our pol-aOC gastruloids also exhibit key aspects of heart morphogenesis, including the formation of internal cavities, endocardial differentiation, and vascular development. scRNA-seq analysis identifies cardiomyocytes with an atrial-like phenotype, and additional features suggest the potential emergence of valve structures, as indicated by SOX2+ cavities, or the expression of valve primordia markers. Besides, the cardiac part in our system co-developed with a vascular network, an endodermal component, and neural lineages, all required to influence cardiac development^47,48^.

## ACKNOWLEDGEMENTS

VV acknowledges support from MBI core funding, CALIPSO grant NRF2019-THE002-0007, and Chaire d’Excellence AMIDEX 2ORGASRH. JBS and RG acknowledge travelling support from CALIPSO grant NRF2019-THE002-0007. JZL acknowledges support from MBI core funding. We thank the Brivanlou’s lab (The Rockefeller University) for their kind gift of the RUES-GLR cell line. We thank MBI microscopy core and MBI microfabrication core for their help and expertise shared.

## AUTHORS CONTRIBUTIONS

MFM, GS and VV designed the research. MFM and GS performed most of the biological experiments and image analysis. AA and GS engineered the aOC system. FD performed 2D adhesion and migration experiments. SS, AM and FL performed image analysis. SBMR and GG designed and made the micro-wells for the soSPIM live imaging experiments. JZ performed bioinformatic analysis for all the single-cell RNA-seq data and LC helped in scRNA-seq data analysis. JBS and RG provided support for live imaging. MFM, GS and VV wrote the manuscript.

## MATERIAL AND METHODS

### Antibodies and reagents

See table S1 for the cell lines, table S2 for antibodies and table S3 for the small molecules and recombinant proteins used in this study.

### Embryonic stem cell culture

We used three human embryonic stem cell (hESC) lines in this study: H1 cell line, H9 cell line and RUES2-GLR cell line, containing three independent fate reporters (SOX2-mCit, BRA-mCer and SOX17- tdTom). H1 cell line was obtained from the group of Yi-Chin Toh (Queensland University of Technology) and RUES2-GLR cell line was obtained as a gift from the group of A. Brivanlou (Rockfeller University). H1 cells were routinely cultured on six-well plates coated with hESC-qualified Matrigel (Corning) in mTeSR media (Stemcell Technologies, Vancouver, Canada) with daily media replacement. The cells were passaged using ReLeSR (Stemcell Technologies) as per manufacturer’s instructions. RUES2- GLR cells were cultured on hESC-qualified, reduced growth factor Geltrex-coated (1:40 dilution, Thermofischer Scientific, Waltham, MA USA) six-well plates and maintained in HUESM medium conditioned by mouse embryonic fibroblasts and supplemented with 20ng/mL bFGF as previously described^49,50^. Cells were passaged using Dispase (Corning) and mechanical scrapping under a microscope to select only the undifferentiated colonies. Both cell lines were cultured at 37°C and 5% CO2.

### Artificial organizing centers (aOCs) preparation and growth factors coating

15 μm streptavidin microspheres (Bangs Laboratories) were resuspended in low-binding microtubes (Eppendorf) in PBS and washed 2 times, before incubation with 100ug/mL biotinylated-heparin (Sigma-Aldrich) for 2 hours at room temperature on a tube rotator. Beads were centrifuged and washed in PBS and subsequently incubated with a cocktail of recombinant growth factors (Activin A 100 ng/mL, BMP4 25 ng/mL, FGF2 10 ng/mL, Wnt3a 100ng/mL). Incubation was done O/N (or up to 48 hours) at 4°C on an orbital shaker. The next day, beads were washed and coated with growth factor reduced-Matrigel for 1 hour at room temperature before gastruloids formation. For gastruloids formation, aOCs were counted and 50 aOCs/gastruloid was used, unless otherwise stated. For pol-aOC gastruloids, the appropriate number of aOCs is seeded in each well of a 24-well AggreWell plate (Stemcell Technologies), previously passivated using Anti-adherence rinsing solution according to the manufacturer’s instructions (Stemcell Technologies). Plates were centrifuged at 100g for 3 minutes to capture aOCs inside the micro-wells before cell seeding.

### Gastruloid formation and culture

Single hESCs were collected from 6-well plates using ReLeSR or Accutase (Stemcell Technologies) and centrifuged. Cells were resuspended in fresh and warm EB formation medium (Stemcell Technologies) containing 10 μM ROCK-inhibitor Y-27632 (Stemcell Technologies) and cell concentration was calculated. The number of cells required to form gastruloids with 150 cells was determined and seeded in each well of a 24-well AggreWell plate (Stemcell Technologies), previously passivated using anti-adherence rinsing solution according to the manufacturer’s instructions (Stemcell Technologies). Plates were centrifuged again at 100g for 3 minutes before storage in the incubator. For most of the experiments, culture medium was carefully changed every 24 h, using a half-replacement method (removing 1 mL of the well, and adding 1 mL of fresh media) until the end of the experiment, at 144 haa. For generation of day 12 samples, gastruloids were transferred to 24-wells ultra-low attachment plates (Corning) at day 8, and media was replaced every 48 h before fixation.

### Treatment of gastruloids with inhibitors/ enzymes

For ECM inhibition experiments, we used 10 mM β-APN (Sigma-Aldrich), 40 μM RGDS peptides (Tocris) and 0.1 mg/mL collagenase IV (Thermofischer Scientific). Working concentrations were set according to previous studies using 3D models^38,51^. Gastruloids were treated from 48 h post-seeding. β -APN and RGDS were diluted in EB formation medium and replaced every day. Collagenase was diluted in EB formation medium, added for 6 hours before washing the cells with fresh and warm medium. This procedure was repeated every day until 144 h post-aggregation, before gastruloids fixation.

### Immunofluorescence and imaging

Gastruloids at different stages after aggregation were collected in 1,5 mL microtubes and fixed 20 min at room temperature in 4% paraformaldehyde and stored at 4 degrees. For immunostainings, gastruloids were transferred to U-bottom 96-well plates or 0.5 mL microtubes and permeabilized in 0.5% Triton-X-100 in PBS for 30 min at room temperature. Gastruloids were then incubated O/N with primary antibodies in 0.2% Triton-X-100 in PBS at 4 degrees while shaking. The day after, gastruloids were washed 3 times (10 min each) with 0.2% Triton-X-100 in PBS and incubated O/N at 4 degrees, or 2 hours at room temperature with secondary antibodies and 2 mg/mL DAPI (Sigma-Aldrich) in 0.2% Triton-X-100 in PBS, while shaking. The day after, gastruloids were washed 3 times in PBS and mounted under a coverslip on glass-bottom petridishes (Ibidi) with RapiClear 1.52 (Sunjin Lab) to clear the samples. The primary and secondary antibodies used in this study are listed in table S2. Confocal images were generated using a spinning-disk confocal microscope (Yokogawa CSU-W1) mounted on a Nikon Eclipse Ti-E inverted body. 405, 488, 561 and 640 nm laser lines were used to excite the different used fluorescent dyes through a Plan Fluor 40X oil immersion lens (NA 1.3, Nikon). A 2x Photometrics Prime 95b CMOS camera was used. Data capture was carried out using Metamorph (Molecular Devices) and Z-stacks were acquired with a Z-step of 1-2 μm. Images were processed using the ImageJ package Fiji.

### 2D adhesion and migration experiments

For 2D adhesion and migration experiments, H1 cells were seeded on top of Matrigel-coated 6-well plates and kept for 72 h in mTeSR media before differentiation using the STEMdiff Definitive Endoderm kit (Stemcell Technologies). Control stem cells were kept in STEMdiff APEL 2 medium (Stemcell Technologies). At 120 h post-seeding, cells were dissociated using Accutase and seeded in 24-well plates (10k cells/well for individual cell spreading and 50k cells/well for cell migration). The wells were previously coated with 1X Matrigel, 10 μg/mL fibronectin (Cytoskeleton), 20 μg/mL collagen I (rat tail, Corning) or 10 μg/mL laminin (Cytoskeleton). Cells were cultured for 4 hours at 37 degrees on the different substrates before fixation or imaging. Cell nuclei were stained with DAPI and cytoskeleton with phalloidin-488, which was used to evaluate cell number and spreading area after segmentation. For migration experiments, live imaging was performed using widefield microscopy and a 10X objective as previously described^52^. Phase-contrast images were acquired at intervals of 4 min over a duration of 12 h. Migration speeds of individual cells and confinement ratios were assessed using the TrackMate plugin in Image J.

### RNA extraction and RNA-seq libraries preparation and sequencing

Gastruloids at 48, 144 haa and after 12 days of culture were collected, washed in PBS before dissociation with Accutase treatment. Samples were all FACS-sorted to select only the cells and discard the aOCs/beads. The experiments were done with the services of NovogeneAIT Genomics. They followed the standard protocol from 10X Genomics Chromium Single Cell 3’ (v3.1 Chemistry dual index). Librairies for single cell gene expression profiling were constructed with 10X Genomics Chromium v3.1 Chemistry Dual Index and sequenced with NovaSeq6000, PE150, 120G of raw data per sample. Sequencing reads were processed with CellRanger 7.1.0. Reads were aligned to the human genome GRCh38-2020-A from 10X Genomics. The raw count matrices were imported in R v4.4.1 with Seurat 5.2.1. Low quality cells were filtered out with different thresholds on gene expression and mitochondrial content. The human gastrulating embryo dataset from Tyser et al, 2020 was used as a reference database for annotating the different clusters. Extraembryonic tissues were excluded from the database since they are not present in gastruloids. We alternatively used CASSIA tool (Xie et al, biorxiv 2024) for annotating the unknown clusters, especially in the day 12 gastruloids. We used ScType to do a first estimation of cell types present in the gastruloids. Doublet detection was carried out with the R package DoubletFinder v1.20.0. The alignment was confirmed to be biologically meaningful by manually verifying that highly variable genes and canonical marker genes were overall well matched with the annotations. We then computed an integrated UMAP for 48-144 h samples as well as iso-aOCs and pola-aOCs gastruloids. At day 12, a close-up view of the cardiomyocyte cluster was generated by recomputing annotations using a human heart atlas (De Bono et al, Development 2025). Differentially expressed genes (DEG) were determined with the Seurat function FindAllMarkers. The detection compares each cluster with the rest of the data.

### Live imaging experiments

JeWells were prepared as already described^20^. Microwells were treated with 0,5% w/v lipidure (Amsbio, CM5206) in 100% ethanol, to prevent the samples from attaching to the wells. The remaining lipidure solution was removed and the wells were dried O/N at room temperature until the residual ethanol evaporated. The next day, JeWells were washed with PBS and EB formation medium and degassed for the liquid to fill up the microwells. Gastruloids were formed as described in the methods section. At 24 haa, gastruloids were harvested using wide-bore tips, transferred in 1,5 mL microtubes and centrifuged. Samples were then transferred to JeWells mounted in 35 mm ibidi Petri dishes and placed in the incubator for 5 min to allow proper seeding. After 5 min, the wells were rinsed to remove excess gastruloids and fresh medium was replaced using EB formation medium, ROCK inhibitor and HEPES. Gastruloids were imaged live using the soSPIM system^19,20^ at 37 degrees, 5% CO2. Data capture was carried out using the Metamorph software. We used a 20X Apochromat Nikon objective. YFP and GFP were sequentially excited with 488 and 555 nm lasers. A volume of 200 μm was acquired with a Z-step of 2 μm between slices, and pictures were captured every 15 min for 72 hours.

### Determination of the proportion of cell lineages from 3D images

Data analysis was performed on maximum intensity projection images. Briefly, maximum intensity projections were generated, before segmentation of each channel (DAPI, SOX2, T/Bra, SOX17) using threshold-based intensities. A binary mask was created for each channel, and the number of positive pixels was calculated using the Analyze Particles function in Fiji. Manual correction was applied to the masks when needed (for example, when 2 gastruloids were in close contact). Normalization was applied by dividing the number of positive pixels for each channel by the gastruloids total area, as determined by the number of pixels in the DAPI channel. For gastruloid size quantifications, the DAPI channel was used to calculate the surface area of gastruloids.

### Centroid maps generation

The different cell types were segmented using threshold-based intensity on maximum intensity projection images, as described above. Gastruloids were oriented according to their longest axis (using the DAPI channel) corresponding to the polarization axis for polarized gastruloids. The position of the centroids for each channel was then calculated and displayed to create a spatial probability density map (see Figure 1E). This routine was done in Python using a custom-made script.

### Proximity of SOX17+ cells with fibronectin fibers

To quantify the presence of fibronectin tracks in gastruloids, we reviewed maximum intensity projection images of n=65 gastruloids spanned across N=10 distinct experiments and defined a track as a fibronectin fiber measuring at least 20 μm and present in the SOX2^+^ area, opposite from the SOX17^+^ region. To assess the proximity of those tracks with SOX17^+^ cells, we used maximum intensity projections images of 5-10 slices (equivalent to 10-20 μm, approximately the size of a cell) around a track, and assessed the presence of a SOX17^+^ cell in this region.

### Number of SOX17+ poles upon RGD treatment

To evaluate the number of SOX17+ poles upon RGD stimulation, maximum intensity projections of the SOX17 channel was done, before generation of a binary mask based on threshold intensity. We used the Analyze Particle plugin and counted the objects above 20,000 pixels^2^, corresponding to the size of a cluster of SOX17^+^ cells.

### Statistics and reproducibility

The number of samples and independent experiments are given in the figure legends. No statistical method was used to predetermine sample size. The experiments were not randomized. The investigators were not blinded to allocation during experiments and outcome assessment.

## EXTENDED DATA FIGURES LEGENDS

**Extended data figure 1:**
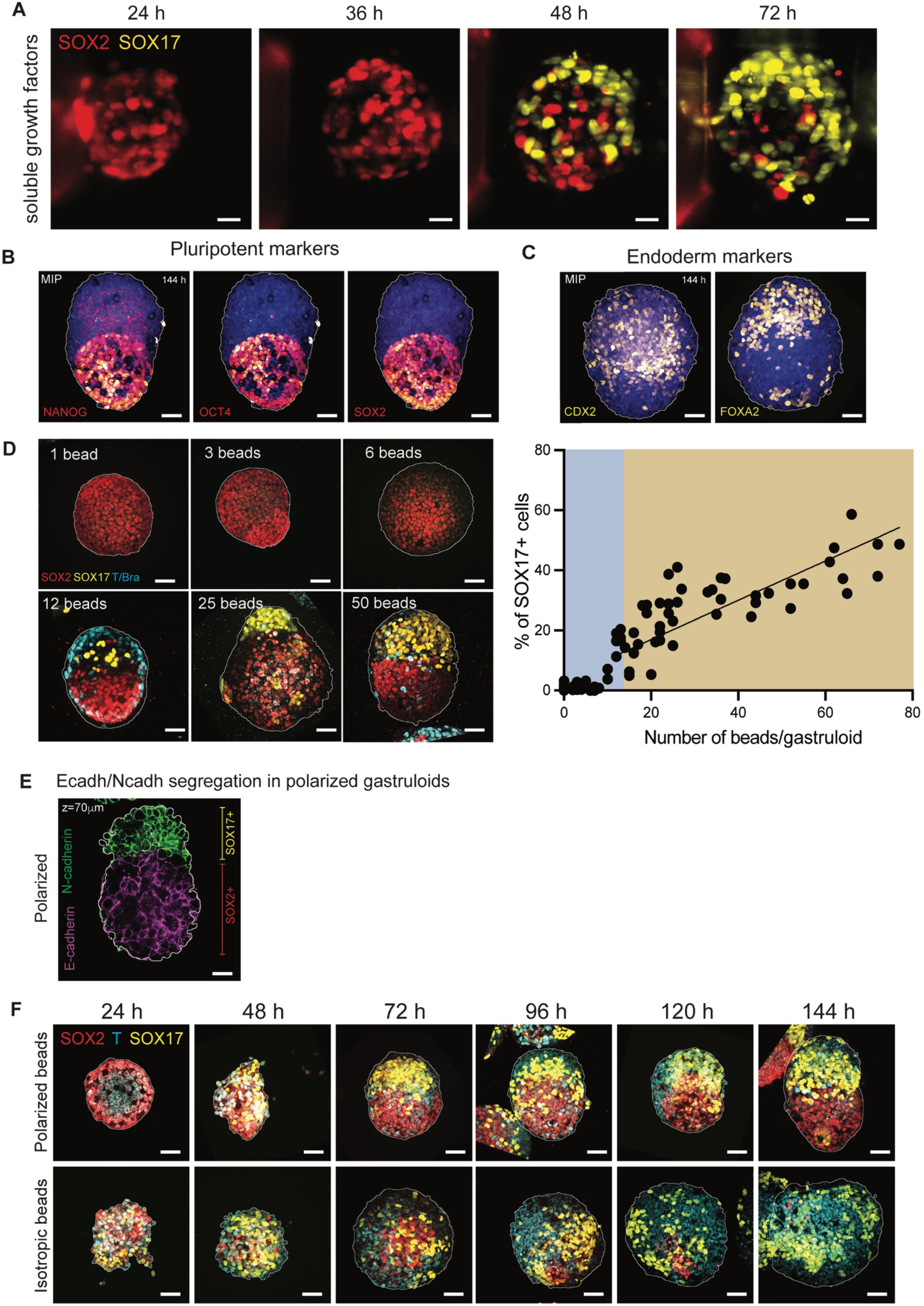
Gastruloids characterization. **(A)** Time-lapse imaging of a single gastruloid differentiated upon soluble growth factors addition in the culture media at 24 h after aggregation. Scale bar: 20 μm. **(B)** and **(C)** Polarized gastruloids were stained with pluripotent (NANOG, OCT4, SOX2) and endoderm (CDX2, FOXA2) markers at 144 h after aggregation. Scale bar: 50 μm. **(D)** Differentiation of gastruloids at 144 h after aggregation as a function of the number of beads included at the beginning of the experiment (1, 3, 6, 12, 25, 50 beads). A graph showing the fraction of SOX17^+^ cells at 144 haa as a function of the number of beads/gastruloid is shown. There is a minimum number of about 10-12 beads needed to trigger differentiation (below this number, the differentiation does not occur: blue area on the graph). A linear regression was performed from x=12 beads, in the brown area of the graph (R^2^ = 0,7753). 5 experiments were analyzed, n= 73 gastruloids. **(E)** E-cadherin/N-cadherin distribution in pol-aOC gastruloids at 144 h after aggregation. Scale bar: 50 μm. **(F)** Data relative to Fig.1G. Representative images of SOX2, T/Bra and SOX17 distributions in pol-aOC and iso-aOCs gastruloids at 24, 48, 72, 96, 120 and 144 h after aggregation. Scale bar: 50 μm.

**Extended data figure 2:**
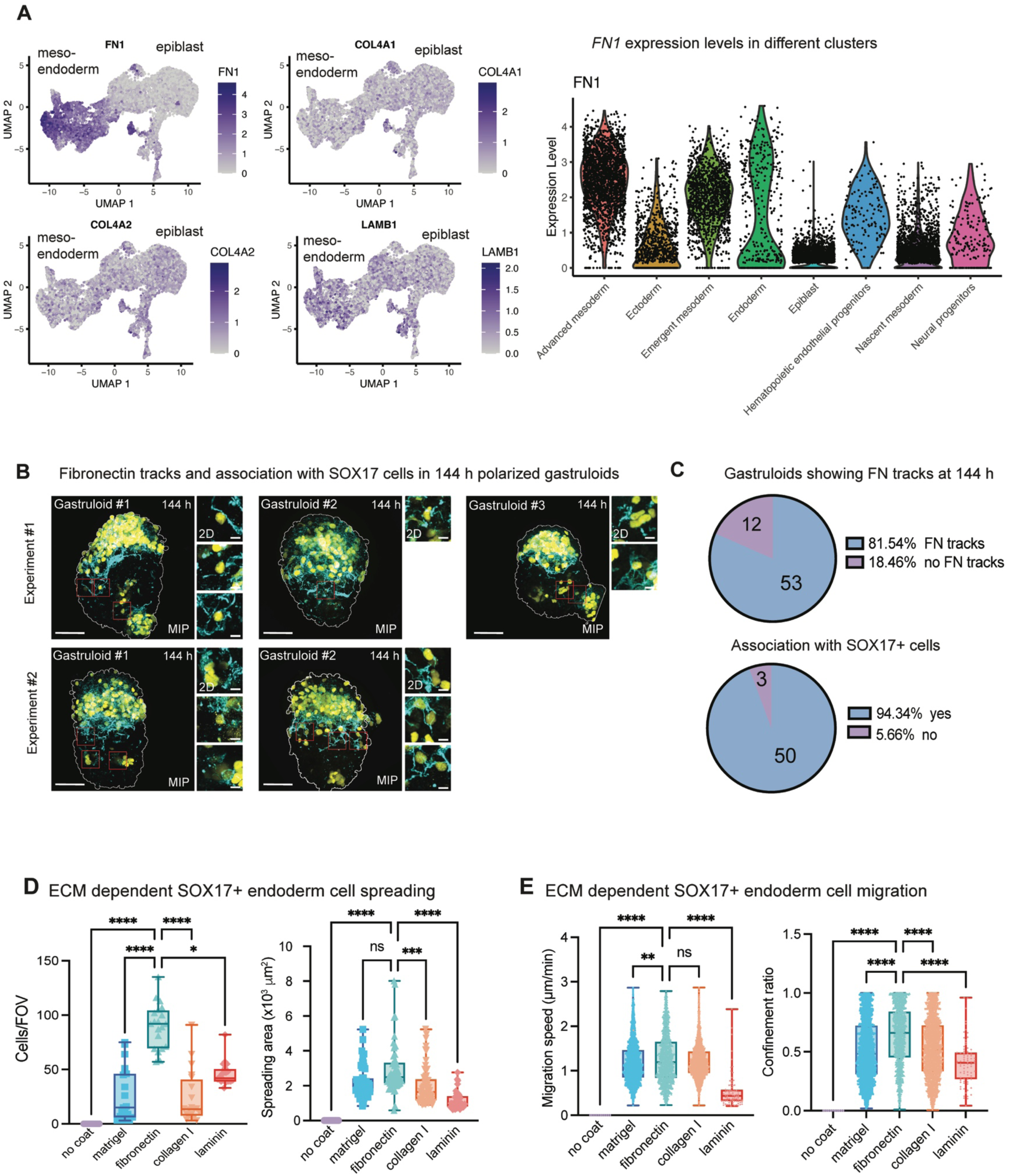
Fibronectin tracks and role in SOX17+ cells migration. **(A)** UMAP plots showing the expression of FN1, COL4A1, COL4A2, LAMB1 in pol-aOC gastruloids. The left part of the UMAP corresponds to meso-endoderm cells whereas the right part represents the epiblast cells. FN1 expression level is also showed in the different cell clusters, showing a higher expression in meso-endoderm clusters as compared to epiblast. **(B)** Examples of fibronectin tracks (cyan) in polarized gastruloids at 144 h in 2 sets of experiments. Large fields of view are maximum intensity projections of the whole volume of the gastruloid, whereas zoomed areas are small maximum intensity projections (max 4-5 planes of 2 μm) to show association of the tracks with SOX17^+^ cells (yellow). Scale bars: 50 μm, zoom: 10 μm. **(C)** Proportion of gastruloids (n=65) showing fibronectin tracks at 144 haa (top row) and proportion of these gastruloids (n=53) showing association or not of these tracks with SOX17^+^ cells (bottom row). **(D)** ECM-dependent cell spreading and **(E)** ECM-dependent cell migration. Cells/FOV are showed, n= 20 fields of view were analyzed for each ECM coating. Statistical comparison made via Kruskall-Wallis test (**** indicates p<0.0001; * indicates p=0.0313). Spreading area (μm^2^) is showed for each of the ECM coatings and was calculated from F-actin/phalloidin stainings. Sample size is n=0 cells (no coat); n=56 cells (Matrigel); n=57 cells (fibronectin); n=61 cells (collagen I); n=46 cells (laminin). Statistical comparison made via Kruskall-Wallis test (ns indicates p=0.0607; *** indicates p= 0.0003; **** indicates p<0.0001). For migration speeds, sample size is n=1834 analyzed tracks for Matrigel, n=1379 analyzed tracks for fibronectin, n=2414 analyzed tracks for collagen I, n=79 analyzed tracks for laminin. Statistical comparison made via Kruskall-Wallis test (ns indicates p>0.9999; ** indicates p=0.0048; **** indicates p<0.0001). Confinement ratio was calculated to assess migration efficiency (1= migration is straight from one point to another; 0 = migration is random and non-directed). Sample size is identical to the previous graph. Statistical comparison made via Kruskall-Wallis test (**** indicates p<0.0001). For **(D)** and **(E)**, Minimum-to-maximum box plots show the 75th, 50th, 25th percentiles

**Extended data figure 3:**
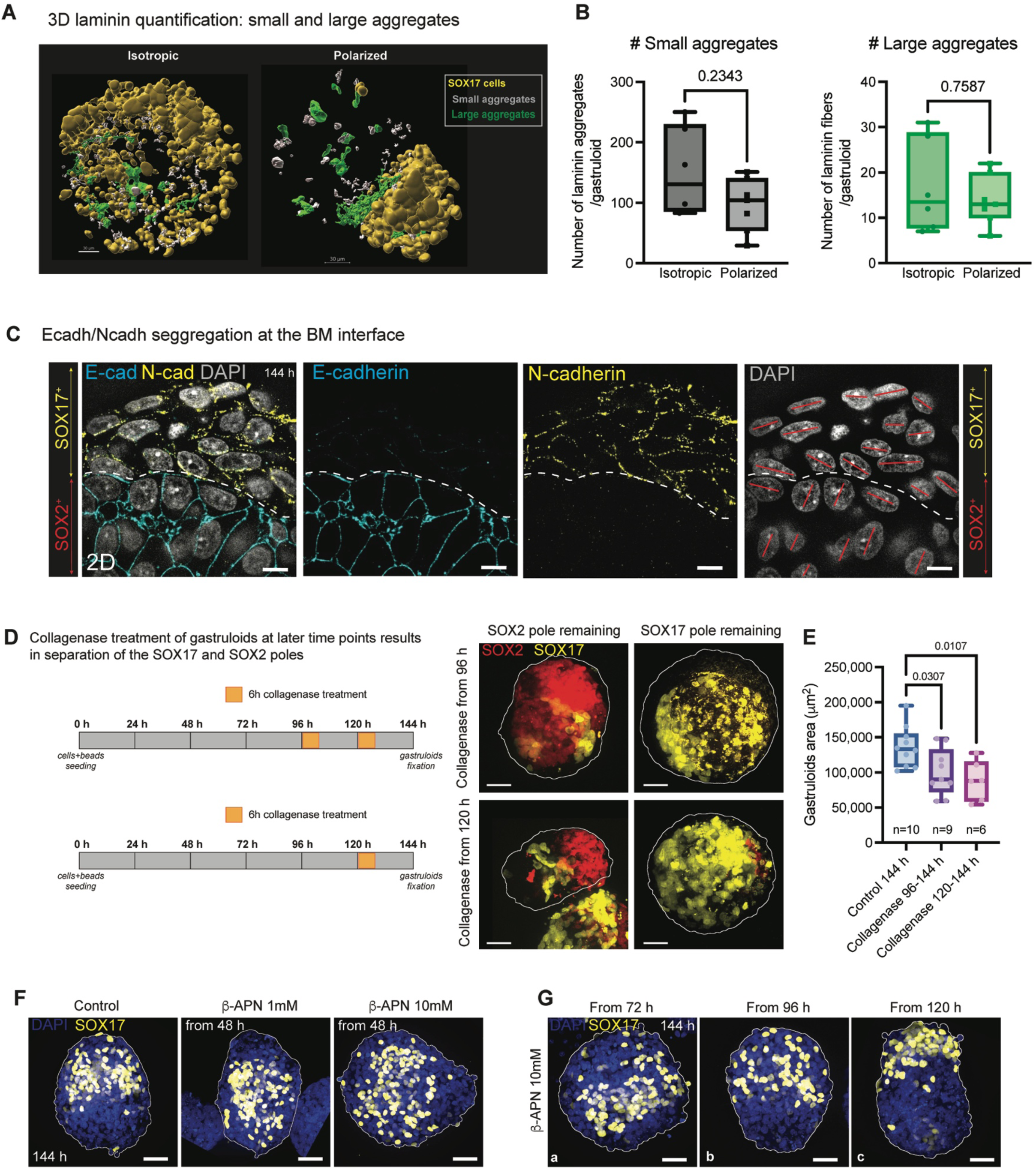
Collagen IV and laminin deposition and maintenance of the stem cell niche. **(A)** 3D reconstructions showing laminin small (grey) and large (green) aggregates in iso-aOC and pol-aOC gastruloids. **(B)** Number of small and large aggregates/gastruloid is showed (n=6 isotropic gastruloids and n=7 polarized gastruloids). Statistical comparison made via Mann-Whitney tests (ns indicates p=0.2343 for aggregates and p=0.7587 for fibers). **(C)** Nuclei orientation in the SOX17+ area (cells expressing N-cadherin) and in the SOX2+ area (cells expressing E-cadherin). Scale bar: 10 μm. **(D)** Experimental procedure for collagenase (100μg/mL) treatment of gastruloids from 96 haa and 120 h after aggregation. Representative images for SOX2 and SOX17 stainings of gastruloids treated from 96 or 120 haa. In both cases, 2 phenotypes were observed, either the SOX2 pole is remaining (left column) or the SOX17 pole is remaining (right column). Scale bar: 50 μm. **(E)** Gastruloid area (μm^2^) is showed for control (n=10 gastruloids) and collagenase-treated gastruloids from 96 haa (n=9 gastruloids) or 120 h after aggregation (n=6 gastruloids). Statistical analysis made via ordinary one-way ANOVA (* indicates p=0.0307 for 96 haa and p=0.0107 for 120 haa). **(F)** Dose-response experiment of β-APN treatment (1 mM every day from 48 haa or 10 mM every day from 48 haa; gastruloids were fixed at 144 haa). Scale bar: 50 μm. **(G)** Time-line experiment of 10 mM β -APN treatment. β -APN was added from 72 haa (a), from 96 haa (b) or from 120 haa (c). Scale bar: 50 μm.

**Extended data figure 4:**
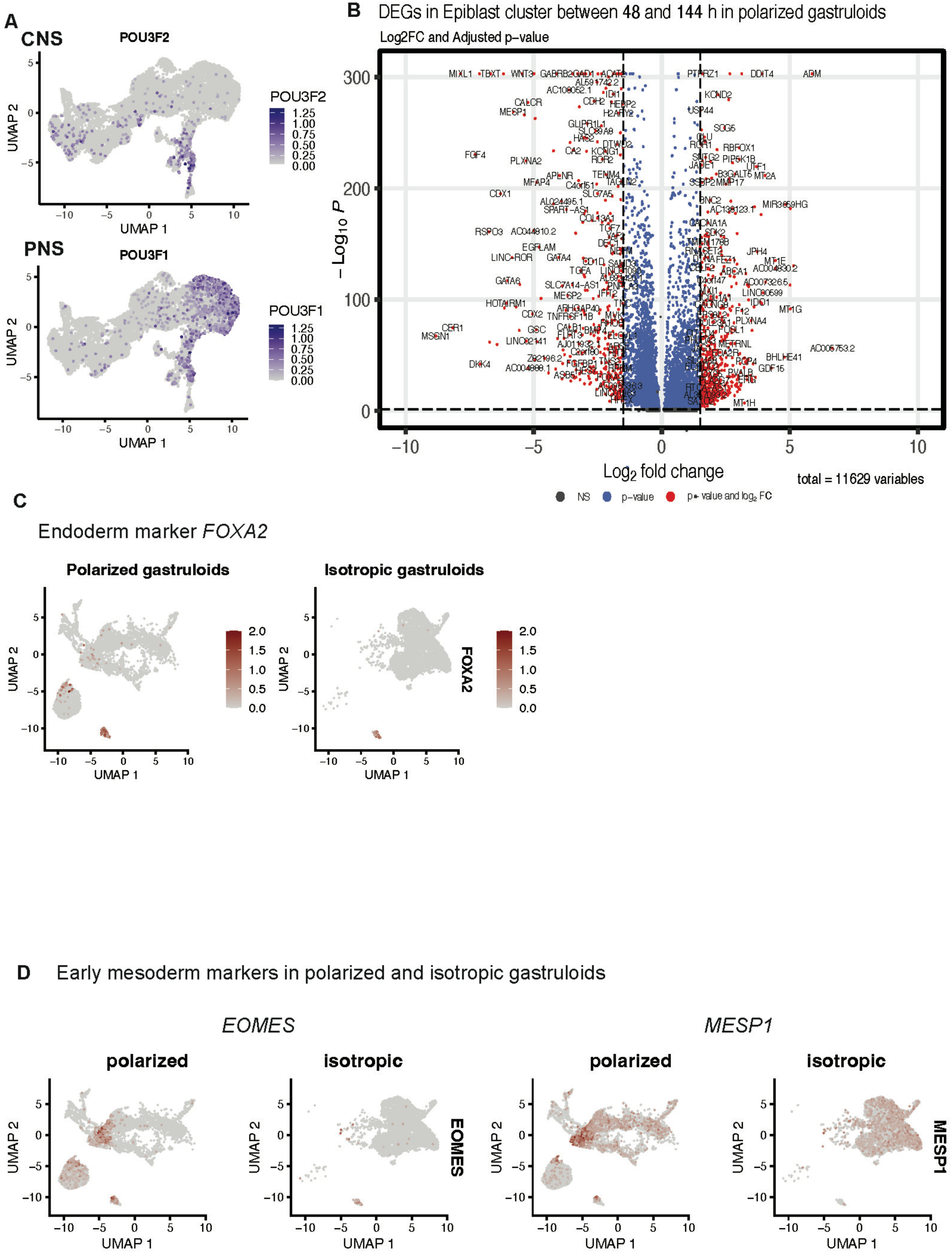
scRNA sequencing of polarized and isotropic gastruloids. **(A)** UMAP plots showing the expression of PNS and CNS markers (*POU3F1* and *POU3F2*) in polarized gastruloids **(B)** Volcano-plot showing the differentially expressed genes in both epiblast clusters between 48 and 144 haa gastruloids. **(C)** UMAP plots showing the expression of endoderm markers *SOX17* and *FOXA2* in polarized and isotropic gastruloids at 144 haa. **(D)** UMAP plots showing the expression of early mesoderm markers in polarized and isotropic gastruloids.

**Extended data figure 5:**
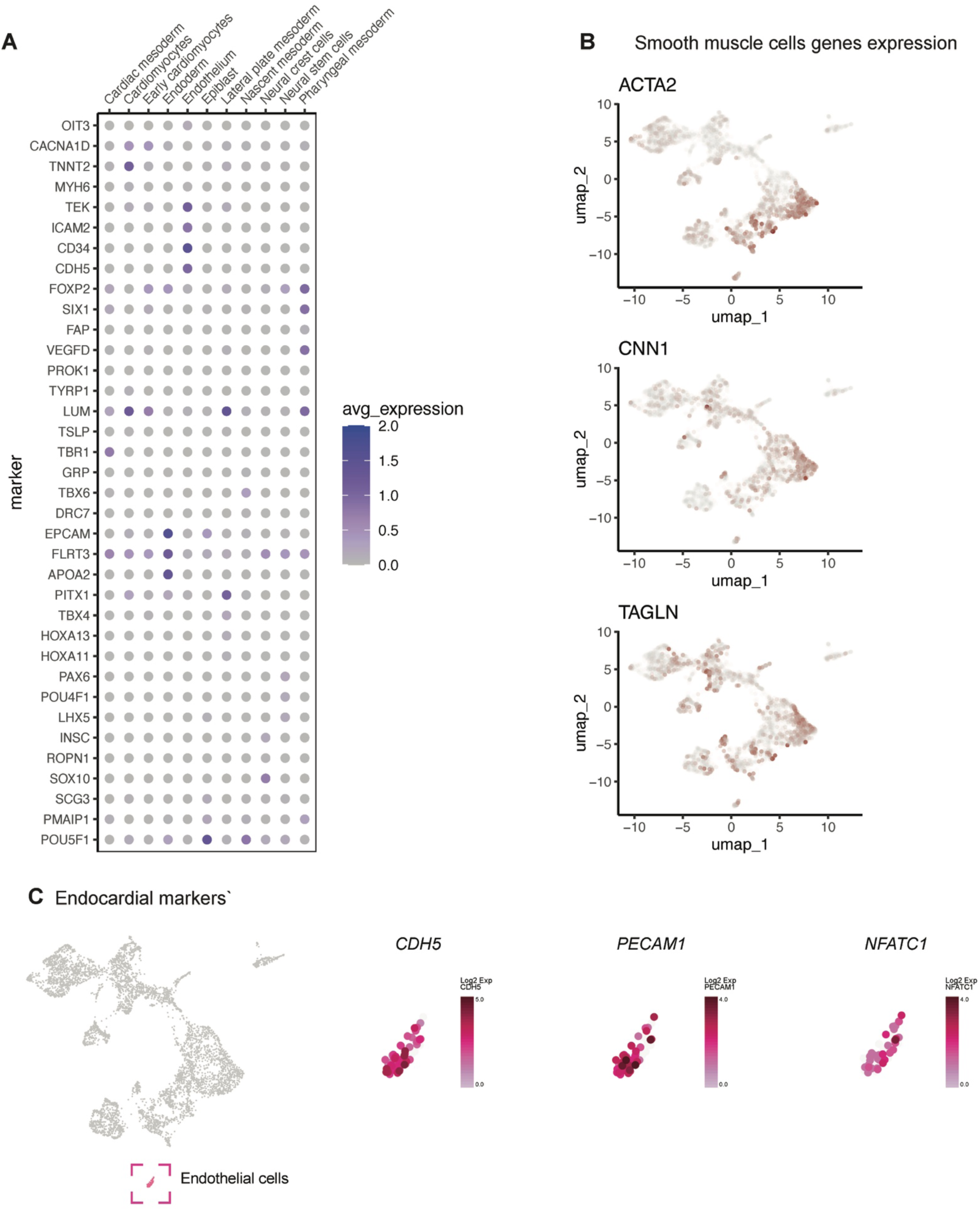
Cardiovascular features in scRNA-seq data. **(A)** Heatmap showing the expression of different genes in the different clusters in day 12 pol-aOCs gastruloids. **(B)** UMAP plots showing some smooth muscle cells gene markers and endocardial markers **(C)** in day 12 gastruloids.

